# Systematic assessment of next generation sequencing for quantitative small RNA profiling: a multiple protocol study across multiple laboratories

**DOI:** 10.1101/113050

**Authors:** MD Giraldez, RM Spengler, A Etheridge, PM Godoy, AJ Barczak, S Srinivasan, PL De Hoff, K Tanriverdi, A Courtright, S Lu, J Khoory, R Rubio, D Baxter, TAP Driedonks, HPJ Buermans, ENM Nolte-‘t Hoen, H Jiang, K Wang, I Ghiran, Y Wang, K Van Keuren-Jensen, JE Freedman, PG Woodruff, LC Laurent, DJ Erle, DJ Galas, M Tewari

**Author notes:** These authors contributed equally (MD Giraldez, RM Spengler).

## Abstract

Small RNA-seq is increasingly being used for profiling of small RNAs. Quantitative characteristics of long RNA-seq have been extensively described, but small RNA-seq involves fundamentally different methods for library preparation, with distinct protocols and technical variations that have not been fully and systematically studied. We report here the results of a study using common references (synthetic RNA pools of defined composition, as well as plasma-derived RNA) to evaluate the accuracy, reproducibility and bias of small RNA-seq library preparation for five distinct protocols and across nine different laboratories. We observed protocol-specific and sequence-specific bias, which was ameliorated using adapters for ligation with randomized end-nucleotides, and computational correction factors. Despite this technical bias, relative quantification using small RNA-seq was remarkably accurate and reproducible, even across multiple laboratories using different methods. These results provide strong evidence for the feasibility of reproducible cross-laboratory small RNA-seq studies, even those involving analysis of data generated using different protocols.

## (Introduction without separate heading below)

RNA-seq using next generation sequencing has been a transformative technology that has been widely used as a method for characterizing the transcriptome in a wide range of biological contexts^1, 2^. Applications of RNA-seq fall into two categories: long RNA-seq and small RNA-seq, distinguished not only by the size of the targeted RNAs, but also by the technical methods used and the resulting biases of the different approaches in the quantitative data produced^3^. For example, the production of the libraries for long RNA-seq, by virtue of having sufficiently long target RNA lengths, commonly utilizes primers (e.g., random primers or oligo-dT) for direct generation of cDNA from RNA. However, small RNA-seq library construction methods typically require an RNA ligation or polyA tailing step to overcome the challenge of performing reverse transcription and subsequent PCR-based amplification from extremely short (e.g., 16-30 nt) target RNA sequences.

Multiple approaches have been developed to overcome the challenge of uniformly and robustly generating cDNA from small RNAs for the purpose of small RNA-seq library preparation^4–9^. Protocols in use for small RNA-seq therefore vary more widely than those used for long RNA-seq, creating significantly more potential for variation across small RNA-seq results using different library preparation protocols and by different labs. In addition, small RNA-seq is increasingly used to study small RNAs present in very low input concentration samples (e.g., in exosomes and other types of extracellular vesicles (EV)^10–19^, or in RNA-protein complexes present in biofluids^20–26^). Normalization methods^27–29^ developed to correct for variation in long RNA-seq data are typically not well-suited for such small RNA-seq data, making it even more important to understand the intrinsic performance characteristics of small RNA-seq methodologies. Whereas performance characteristics such as reproducibility and quantitative accuracy have been well-studied for long RNA-seq^30, 31^, further systematic analyses for small RNA-seq are needed, especially across multiple library protocols using well-defined common reference samples. Yet, with the rapid accumulation of small RNA-seq data (e.g., NIH short-reads archive^32, 33^, EV-associated small RNA sequencing databases^34–36^, TCGA^37^, the exRNA Atlas,^38^ etc.), meaningful, quantitative interpretation of results, especially across different studies, will not be possible without a systematic examination of technical bias, accuracy and reproducibility of small RNA-seq. The analysis of standardized common reference RNA samples, ideally including samples in which “ground truth” of absolute concentrations is known, across multiple laboratories and multiple protocols is critical to such an analysis.

Here, we report a study led by investigators from the NIH-funded Extracellular RNA Communication Consortium (ERCC)^39^ involving nine laboratories, which performed a systematic multi-protocol, multi-institution assessment of the accuracy, reproducibility and technical bias of small RNA-seq using standardized, common reference reagents (i.e., synthetic small RNA pools of defined composition and concentrations, as well as biologically-derived reference RNA). We also sought to identify experimental and computational strategies to reduce the impact of technical bias and protocol-specific effects, in order to improve the accuracy and cross-platform comparability of small RNA-seq results.

## Results

### Study design and standard reference RNA materials

In order to evaluate the performance of multiple small RNA-seq library preparation protocols across multiple laboratories, we developed a set of standard reference samples for distribution as well as a standardized study design (shown in Figure 1). We constructed and distributed to each lab a detailed set of instructions for library preparation and sequencing, along with four reference RNA samples (***Supplementary Tables 1 and 2***): i) an equimolar pool comprising 1,152 synthetic RNA oligonucleotides, corresponding predominantly to human microRNA sequences, as well as a small proportion of non-microRNA oligonucleotides of varied sequence and length (15-90 nucleotides (nt)); ii) two synthetic small RNA pools, called ratiometric pools A and B, each containing the same 334 synthetic RNAs, but in which subsets of RNAs vary in relative amount between pools A and B by 15 different ratios, ranging from 10:1 to 1:10; and iii) blood plasma RNA isolated from human plasma pooled from 11 individuals.

**Figure 1.**
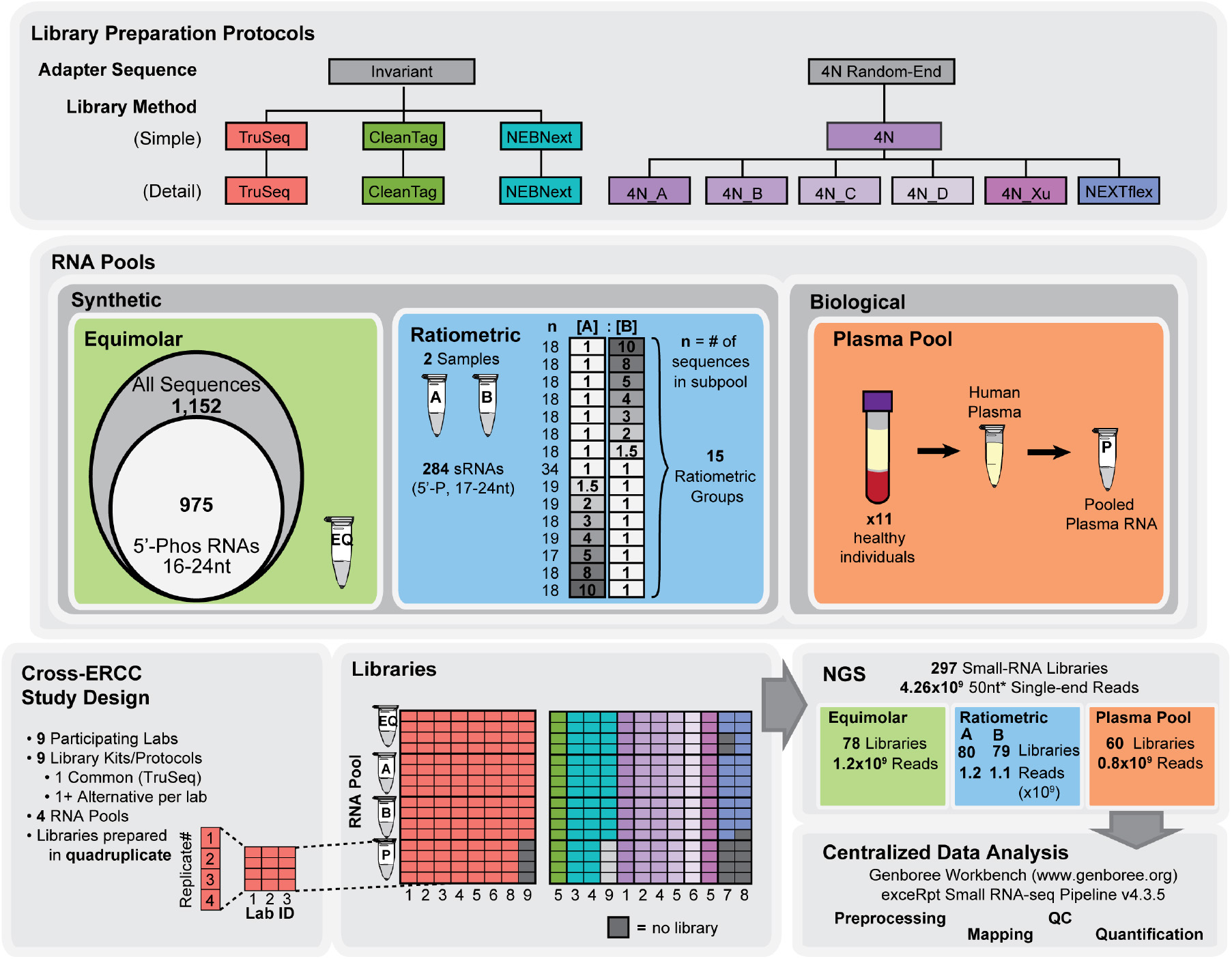
Overview of study design. This figure illustrates the key aspects of this Cross-ERCC study. (**Top**) Nine different library preparation protocols were tested, broadly grouped as “Invariant” or “4N Random-End” according to the absence or presence, respectively, of 4N random terminal bases in their adapter sequences. Three commercially-available kits (TruSeq, CleanTag and NEBNext) with “invariant” adapters were tested. Six 4N Random-End protocols were also tested, and included one commercial kit (NEXTflex), one previously-published protocol (4N_Xu) and four “in-house” protocols (4N_A, _B, _C and _D), developed by ERCC-member labs. The “Simple” and “Detail” boxes introduce nomenclature and color schemes carried throughout the manuscript. (**Middle**) The four RNA pools used in this study as common reference samples are illustrated. The Equimolar and Ratiometric pools were chemically-synthesized to establish “ground-truth” knowledge of absolute and relative abundances, respectively. The Equimolar pool consisted of 1,152 synthetic RNAs (15-90nt) at equimolar concentration. Downstream analyses focused on the subset of 975 RNAs 16-24 nt in length and with 5’-Phosphate modifications. The two Ratiometric Pools, A and B, consisted of 334 synthetic RNAs, in which subsets of the oligos were made to vary in relative abundance between the two pools. The sRNAs were divided into 15 ratiometric subgroups, as shown in the figure, with relative ratios between sample A and B ranging from 1:10 to 10:1. The subset of 284 sRNAs, 17-24 nt in length, were used for downstream analyses, and the number of sRNAs in each ratiometric subgroup is indicated. The Plasma RNA Pool was used to assess performance on low-input biological sample. RNA was centrally isolated and pooled from 11 healthy male individuals and distributed to the participating labs. (**Bottom**) Common reference RNA pools were distributed to 9 Participating labs. Each lab was asked to sequence each of the pools in quadruplicate, using a standardized common protocol (TruSeq) and at least one additional method of their choice. The “Libraries” panel shows the breakdown of the resulting libraries, with the lab IDs indicated in columns, and the replicates and pools shown in rows. The coloring indicates the library prep method used, and follows the coloring schematic in the top “Library Preparation Protocols” panel. NGS sequencing was performed by each lab independently. A total of 297 sRNA libraries were generated, yielding 4.26 billion single-end reads that were subsequently submitted for centralized processing through the exceRpt Small RNA-seq Pipeline. *One lab did a 75nt run and one used the IonTorrent platform which results in reads of varying length.

The common materials were distributed to nine participating research groups (Laurent lab, UCSD; Erle lab, UCSF, Ghiran lab, BIDMC/DFCI; Nolte-’t Hoen lab, UUTR; Freedman lab, UMass; Wang lab, ISB; Galas lab, PNRI; Van Keuren-Jensen lab, TGen and Tewari lab, UMich). Eight groups prepared and sequenced quadruplicate libraries from each of the reference samples using the Illumina TruSeq small RNA kit which is based on ligation of adapters with defined sequences. The TruSeq kit served as a standard reference protocol in this study, allowing for comparison of reproducibility across multiple laboratories. One of the nine groups used a non-Illumina NGS platform (Ion Torrent) and therefore could not generate libraries using the TruSeq protocol. In addition, all nine groups evaluated one or two additional, alternative small RNA library preparation protocols. The alternative protocols were (i) NEBNext (New England Biolabs) - a widely used protocol using adapters with invariant sequences (i.e., no randomized sequence), (ii) CleanTag (Trilink Biotech) - a protocol using adapters with invariant sequences which are chemically modified to reduce formation of adapter dimers, (iii) NEXTflex (Bioo Scientific) - a commercial protocol incorporating adapters with four randomized nucleotides (i.e., 4N) at each ligating end of the adapters in an effort to reduce sequence bias, and (iv) multiple variations of an in-house protocol that also incorporates adapters with 4N randomized ends. We refer to these in-house protocols collectively as “4N” protocols but, they are not identical and must therefore be evaluated separately. Additional details about library preparation protocols and sequencing platforms used in this study are provided in Supplementary Material.

Sequencing data were all centrally analyzed using the ERCC’s exceRpt Small RNA-seq pipeline. This is a publicly available pipeline specifically designed for the analysis of small RNA-seq data (http://genboree.org/theCommons/projects/exrna-tools-may2014/wiki/Small%20RNA-seq%20Pipeline) which uses its own alignment and quantification engine to map and quantify a range of RNAs represented in small RNAseq data, providing RNA abundance estimates from read counts as well as a variety of quality control metrics (***Supplementary Table 3).*** The nine participating groups collectively contributed 4.26 billion reads, corresponding to sequence data from 297 independent libraries (***Figure 1***). Of these, 286 (96%) satisfied minimum quality criteria (see Sample Filtering section of the Methods for details of QC criteria) and were taken forward for the analyses described below. Unless specifically noted otherwise, in the analysis of equimolar pool libraries, NEXTflex Lab 8 data was excluded due to inadequate read coverage in a majority of libraries. FASTQ files for RNA-seq data, as well as any processed files used in this study are available on GEO at the following link: https://www.ncbi.nlm.nih.gov/geo/query/acc.cgi?token=whipakmajrwprcv&acc=GSE94586

### Characterization of sequence-specific bias of Small RNA-seq protocols

Prior studies of small RNA-seq have revealed that there can be significant sequence-specific bias which appears to originate largely during library preparation^4, 40–42^. However this bias has not been characterized systematically across multiple labs using both standardized and divergent protocols and with common synthetic reference reagents. In this study the equimolar reference pool of 1,152 synthetic RNAs was sequenced by all laboratories. In subsequent data analysis, we sought to examine the RNAs most relevant to the library preparation protocols used. We therefore focused on a subset of 975 RNAs that are 16 - 25 nt in length and 5’-phosphorylated, which are the type of RNAs that the small RNA-seq protocols examined are all designed to sequence. We found that the efficiency of recovery of different RNA sequences varied widely within each protocol examined, confirming that small RNA-seq protocols in use are associated with substantial sequence-dependent bias (***Figure 2a, 2b***). The recovery of equimolar RNA sequences varied by multiple orders of magnitude across sequences, much more so than the bias reported for long RNA-seq^31^. However, the sequence-specific bias in small RNA-seq was highly reproducible, both within a given protocol across technical replicates and across laboratories using the same protocol, as evident in results from the standardized TruSeq Illumina protocol carried out by multiple laboratories (Figure 2a).

**Figure 2.**
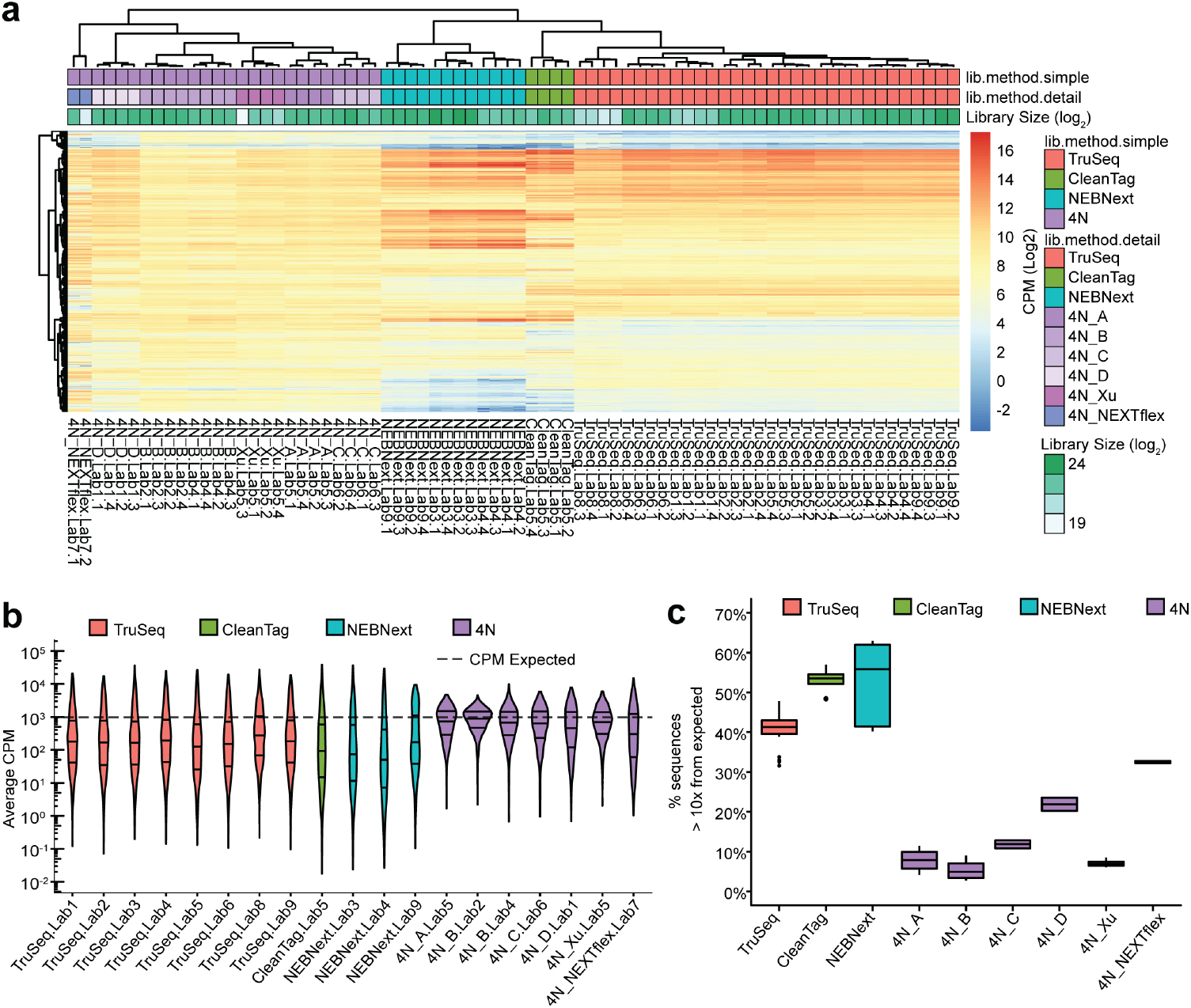
Results from sequencing of the synthetic small RNA equimolar pool by multiple laboratories using multiple library preparation protocols. (**a**) The heatmap shows expression levels for each synthetic RNA sequence (rows) across equimolar pool libraries (columns). Expression levels represent Relative Log Expression (RLE)-normalized, log_2_-scaled CPM calculated for 975 equimolar pool sequences which are 16-245 nt in length and 5’-phosphorylated. Hierarchical clustering for rows and columns represents complete linkage clustering on Euclidean distances (the default setting for the R package, pheatmap, used for plotting). Columns are color-coded (top) to indicate distinct library preparation methods, where “lib.method.simple” indicates the four main method groups (TruSeq, NEBNext, CleanTag and 4N). The “lib.method.detail” annotation coloring additionally indicates the specific in-house or commercial 4N method. Columns are labeled to identify replicate samples. The “Library Size” shaded intensity bar above the columns indicates the total sequencing depth for the individual libraries (log_2_-scaled). (**b**) Violin plots show the mean counts per million (CPM) measured for each detected sequence (y-axis; log10-scaled) across technical replicates from equimolar pool libraries prepared by different institutions and using different library preparation protocols (x-axis). The width of the violins is proportional to the density of data points at each position. The horizontal lines within each violin represent the 25, 50 and 75^th^ percentiles. The dashed horizontal line shows the CPM expected for an equimolar pool of the 975 sequences selected for analysis (10^6^ / 975 miRs = 1025.6). Violins are grouped and colored by distinct classes of library preparation protocols. (**c**) Boxplots show the percentage of equimolar pool sequences expressed at 10x higher or 10x lower than the expected CPM, for each library preparation protocol. Expected counts were computed as total mapped reads / number of unique miRNAs. The fraction of miRNAs was calculated for each replicate library and the boxplots summarize the findings from replicate libraries from all labs for the protocol indicated. The underlying boxes correspond to the the median and IQR. Whiskers represent the 1st/3rd quartile +/− 1.5 * IQR. Boxplots for CleanTag, 4N_A, 4N_C, 4N_D, and 4N_Xu plot values were calculated from four technical replicates from single labs. The number of values plotted for the remaining labs are as follows: TruSeq (n= 32 = 8 labs x 4 replicates), NEBNext (n= 12 = 3 labs x 4 replicates), 4N_B (n = 8 = 2 labs x 4 replicates) and 4N_NEXTFlex (n = 6 = 1 lab x 4 replicates + 1 lab x 2 replicates).

We observed that the libraries prepared by the different participant labs clustered into four groups (Figure 2a) corresponding to the different types of protocols included in the study (TruSeq, NEBNext, CleanTag and 4N), indicating that different protocols have distinct patterns of sequence-specific bias. Consistent with this result, when we selected the ten most overrepresented and underrepresented sequences for each library preparation protocol, these varied widely between protocols (***Supplementary Figure 1***).

Although all library preparation protocols exhibited bias, protocols using randomized adapters showed less bias, as shown by a smaller variation in read abundance across sequences (Figure 2b). As one measure of this, we calculated the median percentage of sequences with a number of reads (i.e., counts per million) more than ten times above or below the expected value, for each of the protocols. This ranged from 41%-56% for all the protocols using adapters with defined sequences (TruSeq: 41%; CleanTag: 54% and NEBNext: 56%), whereas it ranged from 5%-23% for protocols using adapters with randomized nucleotides (4N_A: 8%; 4N_B: 5%; 4N_C: 12%; 4N_D: 22%, 4N_Xu: 7% and 4N_NEXTflex: 17%) (Figure 2c). Also consistent with a reduction in bias using randomized adapters, the 4N in-house protocols showed fewer missing sequences from the equimolar pool as compared to the other protocols (***Supplementary Table 4***). This was also evident when downsampling of datasets was performed so that the same number of total sequencing reads could be compared across protocols, at varying sequencing depths (***Supplementary Figure 2***).

We also sought to assess the degree to which the small RNA cloning biases are reproducible across labs using the same or different protocols, by examining the rank-order of RNA sequence abundance. To see if the rank-order of miRNA abundance was preserved, we calculated Spearman rank correlations for the equimolar pool counts between labs/protocols. As depicted in the heatmap (***Supplementary Figure 3***) the correlation between lab was very strong when using a common standardized protocol [Lib. method: combined Rho value (top/bottom 2%, n samples): TruSeq: 0.98 and (0.89, 1; n = 8), NEBNext: 0.99 (0.93, 1; n = 4)]. This correlation was fairly strong when comparing results from the in-house 4N protocols, with the caveat that these protocols were not identical across labs and the spread using these data points was less relative to TruSeq because of the reduced bias associated with the 4N protocols [4N: 0.76 (0.15-99; n = 6)]. Comparison across labs using different protocols showed weak correlation, consistent with the differences in sequence-specific biases between protocols discussed above [TruSeq vs NEBNext: 0.60 (0.56, 0.66); TruSeq vs CleanTag: 0.73 (0.71-0.77); TruSeq vs 4N: 0.45 (0.18-0.70); NEBNext vs CleanTag.: 0.71 (0.71-0.72), NEBNext vs 4N: 0.51 (0.28-0.52) and CleanTag vs 4N: 0.50 (0.20-0.62).

Accuracy and cross-protocol concordance of small RNA-Seq protocols for relative quantification In addition to assessing bias in recovering different small RNA sequences relative to each other in the same sample, we investigated the accuracy of relative quantification of the same small RNAs between different samples. In order to do so, we designed two ratiometric pools, SynthA and SynthB, each containing the same 334 synthetic miRNA sequences, but with varying relative abundance of RNA sequences between the two samples (***Supplementary Table 2***). Sequencing of these ratiometric synthetic RNA pools allowed us to compare the observed vs. expected values for fifteen different defined expression ratios (1:10, 1:8, 1:5, 1:4, 1:3, 1:2, 1:1.5, 1:1, 1.5:1, 2:1, 3:1, 4:1, 5:1, 8:1 and 10:1) for each of the different library preparation protocols. All of the protocols tested showed close concordance between observed and expected ratios, clearly distinguishing the 15 different ratios (Figure 3). We performed an additional analysis of the data using standard differential expression analysis to determine the smallest fold-difference in abundance that could be distinguished using small RNA-seq methods. We observed that in most datasets (i.e., for most protocols), the majority of miRNAs present at a fold difference of as little as 1.5-fold between the two samples could be detected as differentially abundant (***Supplementary Figure 4).*** As shown in ***Supplementary Table 5***, all the evaluated protocols were remarkably accurate in this sense, in spite of their inherent sequence-specific bias. It is interesting that even the protocols associated with the highest bias (i.e. protocols using adaptors with defined sequences) were demonstrably accurate in most relative quantifications.

Much as we had done for the synthetic equimolar pool, we also used sequencing data from our ratiometric pools SynthA and SynthB to assess the degree to which small RNA cloning biases are reproducible across labs using the same or different protocols, by examining the rank-order of RNA sequence abundance. Taking a general overview of the Spearman rank correlations results obtained for the SynthA and SynthB samples, the results were essentially the same as for the equimolar pool: the correlation was strong when using the same protocol, but weaker across different protocols (Figure 4c). In contrast, when we analyzed the concordance of the obtained ratios SynthA/SynthB (Figure 4c), we found a very strong correlation between labs not only when using the same protocol, but also across different protocols, further confirming that the relative quantification is not subject to substantial protocol-specific confounding effects [Lib. method: combined Rho value (top/bottom 2%, n samples): TruSeq:0.98 (0.98,0.99, n = 8); 4N: 0.98 (0.96,0.99, n = 8); NEBNext:0.98 (0.98,0.98, n = 3); TruSeq vs NEBNext: 0.96 (0.95,0.97, n = 11); TruSeq vs 4N: 0.97 (0.96, 0.97, n = 16); TruSeq vs CleanTag: 0.95 (0.94, 0.96, n = 9); NEBNext vs 4N: 0.96 (0.94, 0.98, n= 11); NEBNext vs CleanTag: 0.95 (0.94, 0.95, n = 4); 4N vs CleanTag: 0.96 (0.93, 0.96, n = 9)].

**Figure 3.**
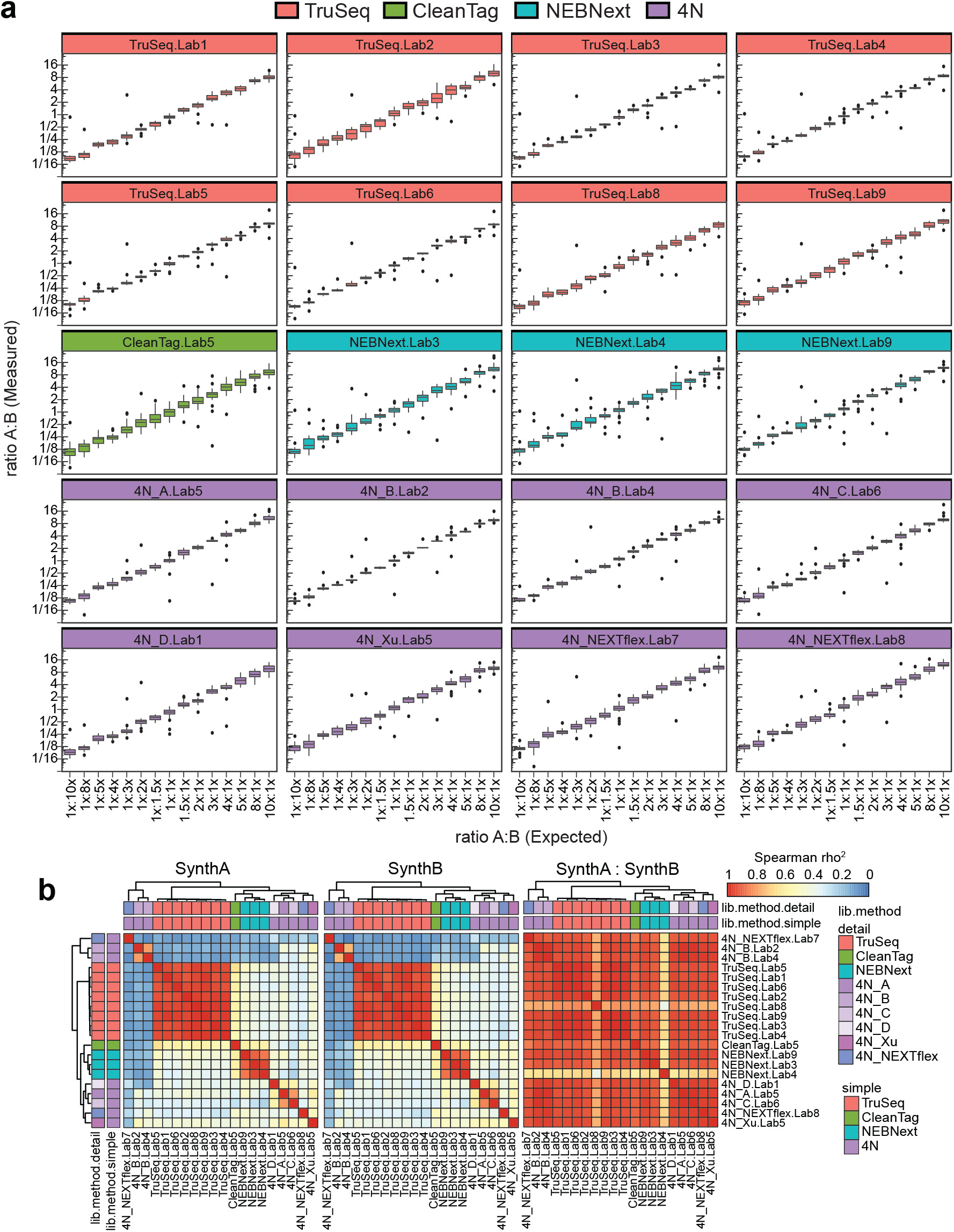
Small RNA-seq accuracy and cross-protocol concordance in measuring relative expression levels between samples. (**a**) Boxplots show the observed ratio (y-axis; log_2_-scale) vs. expected ratio (x-axis) of sets of miRNAs present in the SynthA and SynthB synthetic RNA pools at 15 defined ratios ranging from 1:10 to 10:1 (as listed on the x-axis as “expected” ratios). Separate plots are shown for each sample, which comprises a given library preparation protocol and laboratory. Plots are grouped by library preparation method (TruSeq – pink, CleanTag – green, NEBNext – turquoise and 4N-type protocols – purple). Boxes show the median + IQR; upper/lower whiskers indicate the smallest/largest observation less than or equal to 1^st^/3^rd^ quartile -/+ 1.5 * IQR. (**b**) Heatmaps showing the pairwise, squared Spearman rank correlation coefficients calculated from sequencing the ratiometric synthetic RNA pools A and B. The heatmap to the left (**SynthA**) uses mean CPM data from sequences in the SynthA pool. The heatmap in the middle (**SynthB**) uses mean CPM data for sequences in the SynthB pool. The heatmap to the right (**SynthA: SynthB**) shows Spearman rank correlation coefficients across labs and protocols when using the ratio of the mean CPM for SynthA and SynthB as the input data. Rows and columns are color-coded (top) to indicate distinct library preparation methods, where “lib.method.simple” indicates the four main method groups (TruSeq, NEBNext, CleanTag and 4N). The “lib.method.detail” annotation coloring additionally indicates the specific in-house or commercial 4N method. Hierarchical clustering for rows and columns is the same for all three heatmaps, and is based on the average pairwise Euclidean distances calculated from the SynthA CPM and SynthB CPM correlation matrices.

**Figure 4.**
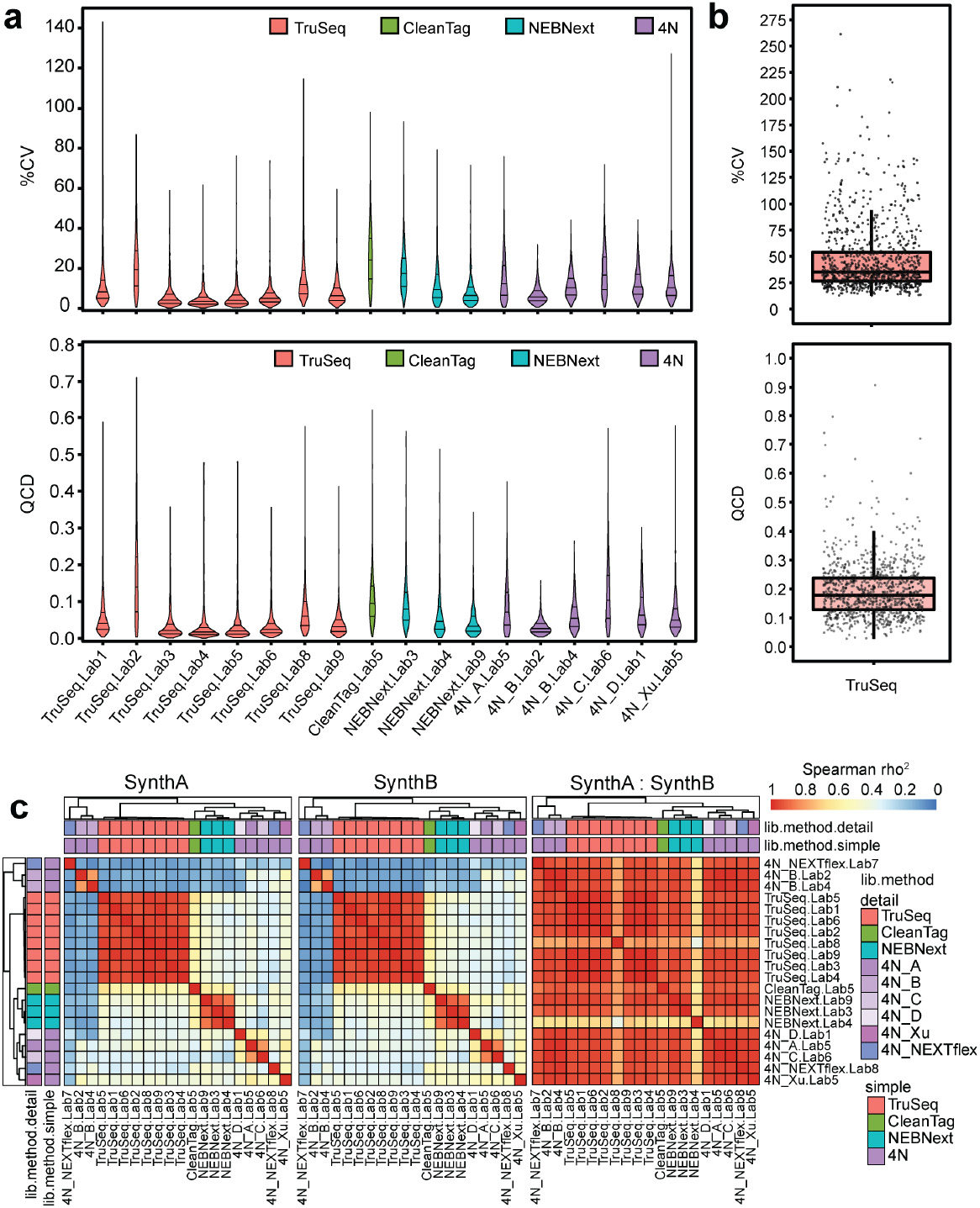
Reproducibility of small RNA-Seq within- and between-labs. (**a**) Violin plots show sequence-wise % Coefficient of Variation (%CV) and quartile coefficient of dispersion (QCD) for equimolar synthetic RNA pool samples, calculated across technical replicates (n = 4 replicates per lab/method). Plots are grouped and color-coded by library preparation method (Truseq – pink; CleanTag – green; NEBNext – turquoise; 4N-type protocols – purple). Horizontal lines within each violin indicate the 1^st^, 2^nd^ and 3^rd^ quartiles. (**b**) Boxplots show between-lab reproducibility of quantification of individual RNA sequences from the equimolar synthetic RNA pool using the TruSeq library preparation protocol. Each dot represents %CV and QCD calculated across all labs for an individual sequence in the equimolar pool. The mean CPM for each sequence across technical replicates was used to calculate the between-lab %CV and QCD plotted here. The overlaid boxes show the median and IQR. Whiskers represent the 1^st^/3^rd^ quartile +/− 1.5 * IQR.

### Reproducibility of small RNA-seq protocols

As each group was asked to contribute four technical replicates for each contributed library preparation protocol, we were able to assess the within-lab reproducibility of each protocol utilized. The heatmap representation of the equimolar synthetic RNA pool sequencing data (Figure 2a) showed that technical replicates of the same protocol clustered tightly together within each lab, consistent with high intra-lab reproducibility of replicates. In order to quantify intra-lab variation from this data, we used two used two metrics, (i) %CV (standard deviation/mean) and, (ii) quartile coefficient of dispersion –QCD- (interquartile range / average of the first and third quartile). The median %CV ranged from 6%-24% for the different protocols. library preparation methods. The %CV for each protocol, written as [Lib. method: Median %CV (top/bottom 2%, n samples where each “sample” is a lab with multiple technical replicates)] were: TruSeq: 6.19% (1.04, 43.52, n = 8); NEBNext: 10.06% (1.59, 51.88, n= 4); 4N-type protocols: 11.22% (1.47, 76.78, n = 6) and CleanTag: 23.69% (3.23, 61.31, n = 1)] (Figure 4a). In addition, the median quartile coefficient of dispersion was <0.1 for all the protocols/labs [Lib. method: Median QCD (top/bottom 2%, n samples where each “sample” is a lab with multiple technical replicates), TruSeq: 0.03 (0.00, 0.30, n = 8); NEBNext: 0.05 (0.01, 0.27, n= 4); 4N-type protocols: 0.06 (0.01, 0.27, n = 6) and CleanTag: 0.09 (0.01, 0.33, n = 1)] (Figure 4a). We also evaluated the intra-lab variation from technical replicates of sequencing the SynthA and SynthB libraries. The calculated %CV and QCD values were similar to those observed for the equimolar libraries (***Supplementary Figure 5***).

In addition to assessing within-lab reproducibility of technical replicates, we were able to characterize reproducibility of small RNA-seq across laboratories because a standardized small RNA-seq protocol (i.e., TruSeq) to be followed by multiple groups was included in the study design. Using the sequencing results for the equimolar pool and treating each laboratory’s results as one trial of the experiment, we calculated the % CV as well as the QCD for the mean CPM values for each RNA sequence across laboratories. The median %CV across labs was < 35.09% (top/bottom 2%: 16.32, 142.75; n = 8) and the median QCD across labs was 0.18 (top/bottom 2%: 0.06, 0.45; n = 8) (Figure 4b). We also calculated across between-lab variation for the SynthA and SynthB pools individually and obtained comparable median %CV and QCD values to those described for the equimolar libraries [Median %CV: SynthA = 30.79%, (top/bottom 2%: 14.50, 66.90; n = 8), SynthB = 32.08% (top/bottom 2%: 14.50, 66.90; n = 8) and Median QCD: SynthA = 0.17 (top/bottom 2%: 0.05, 0.43; n = 8), SynthB = 0.18% (top/bottom 2%: 0.05, 0.43; n = 8).

### Performance of small RNA-seq protocols using biological samples

We also sought to characterize performance of small RNA-seq protocols across labs using standard reference RNA derived from biological material, since factors such as greater complexity of both RNA species and of the biological matrix in which they are present, could produce differing results from those obtained using synthetic reference RNAs. Whereas synthetic reference RNA pools are made to have a defined composition and concentration of RNA species, for biological samples the “ground truth” is not directly known and therefore technical bias and accuracy of small RNA-seq protocols cannot be readily determined. However, sequencing of standardized biological material across labs and protocols can allow evaluation of reproducibility, as well as of diversity of microRNA sequences recovered by different protocols. In order to perform this analysis, we used RNA extracted from a pool of human blood plasma from 11 donors as a standardized reference sample. Aliquots of the extracted RNA were shipped to the participating labs for sequencing in quadruplicate using the same protocols as used for synthetic pools above (i.e., standardized TruSeq protocol as well as one or two alternative protocols run by each lab, unless indicated otherwise). We focused our analysis of sequencing data on microRNAs, since these are a well-characterized class of small RNAs^43^ that have been extensively studied in human plasma^44^.

In order to evaluate the reproducibility of a given protocol within a lab, we analyzed the four plasma technical replicates available within each lab using each particular protocol. In a heatmap clustering analysis of the comprehensive plasma RNA sequencing results (filtered to require a minimum CPM for analysis, as detailed in the legend for Figure 5a), technical replicates of the same protocol clustered together (Figure 5a). We quantified the intra-lab reproducibility of plasma small RNA-seq for each protocol using %CV and QCD calculated for individual miRNA sequences across replicates (Figure 5b). The median %CV across the range of miRNAs analyzed ranged from 9% − 25% for different protocols. This degree of reproducibility was comparable to that observed in small RNAseq of synthetic reference pool RNA using different protocols, as described earlier (Figure 4). More specifically, the median %CV values for sequencing reference plasma RNA by protocol [represented as Lib. method: Median %CV (top/bottom 2%, n samples where each “sample” is a lab with multiple technical replicates)] were: TruSeq: 8.85% (1.64, 33.73, n = 8); NEBNext: 12.09% (3.00, 34.35, n = 4); 4N-type protocols: 10.45% (1.76, 36.18, n = 6) and CleanTag: 25.09% (5.47, 52.33, n = 1)]. In addition, the median QCD was ≤0.1 for all the protocols [represented as Lib. method: Median QCD (top/bottom 2%, n samples where each “sample” is a lab with multiple technical replicates):TruSeq: 0.04 (0.01, 0.17, n = 8); NEBNext: 0.06 (0.01, 0.18, n= 4); 4N-type protocols: 0.05 (0.01, 0.22, n = 6) and CleanTag: 0.10 (0.02, 0.24, n = 1)].

**Figure 5.**
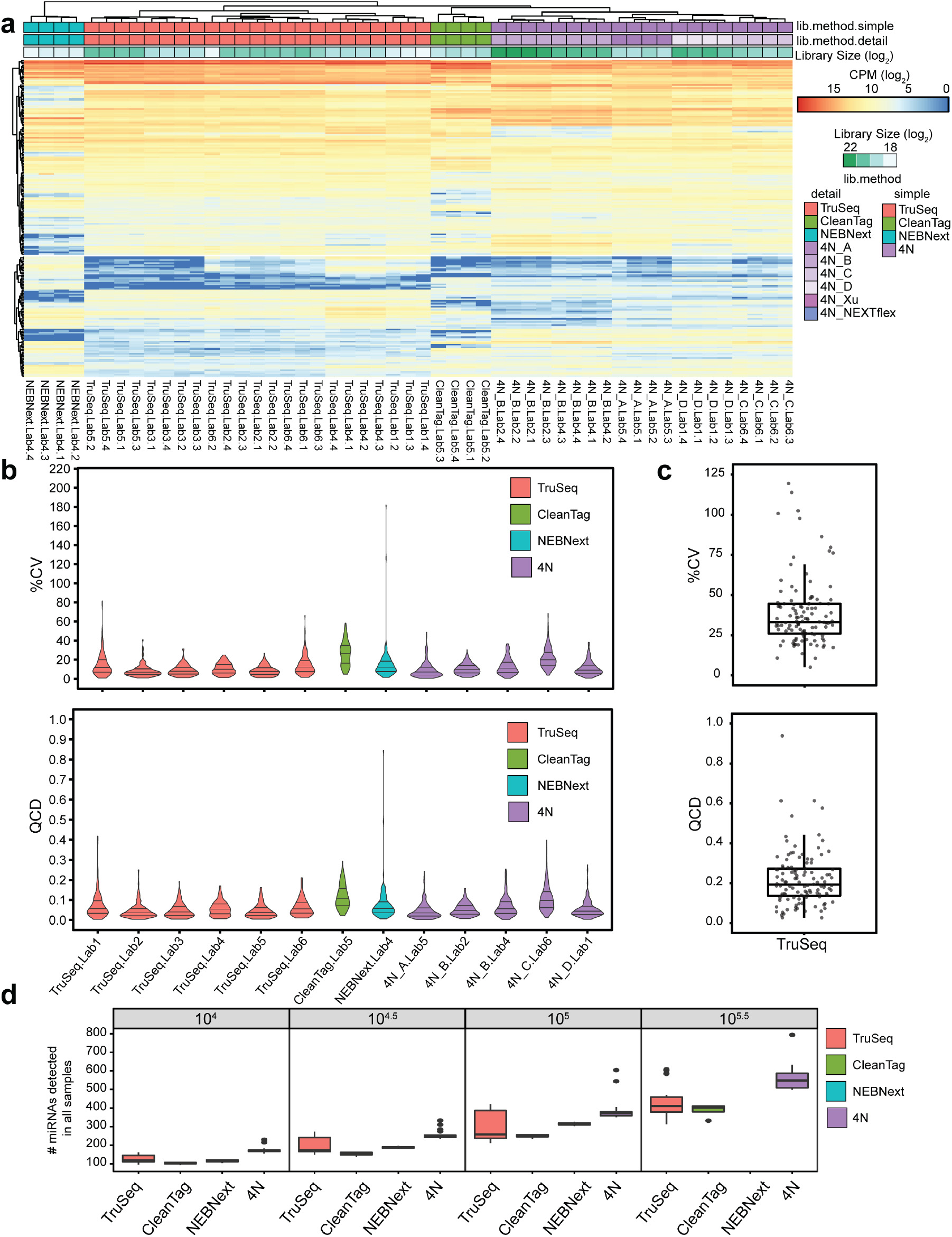
Small RNA-seq of standard reference plasma RNA by multiple laboratories using multiple library preparation protocols. (**a**) The heatmap shows CPM (log_2_ scale) for each sequence (rows) across plasma pool libraries (columns). Only mature miRNAs with a high confidence of detection are shown, requiring a minimum of 100 CPM in 90 percent of samples from at least one protocol (TruSeq, CleanTag, NEBNext or 4N). Hierarchical clustering for rows and columns represents complete linkage clustering on Euclidean distances. Columns are color-coded (top) to indicate different library preparation methods, where “lib.method.simple” indicates the four main method groups (TruSeq, NEBNext, CleanTag and 4N). The “lib.method.detail” annotation coloring additionally indicates the specific in-house or commercial 4N method. Shaded intensity bars on the columns corresponding to “Library Size” indicate the sum of the mature miRNA-mapped read counts prior to filtering for the individual libraries (log_2_-scaled). (**b**) Violin plots show the within-lab %CV and QCD for sequencing of reference plasma RNA technical replicates per each lab/protocol. (**c**) Boxplots show between-lab reproducibility of individual miRNA sequences from sequencing of standard reference plasma RNA using the TruSeq library preparation protocol. Each dot represents %CV or QCD calculated across labs for a different sequence. The between-lab %CV and QCD were calculated using the mean CPMs for each sequence across technical replicates for each lab. The underlying boxes show the median and IQR. Whiskers represent the 1st/3rd quartile +/− 1.5 * IQR. (**d**) Boxplots show the number of mature miRNAs detected by each protocol based on downsampling of datasets to equivalent sequencing depths (see Methods for more details). Each box summarizes number of miRNAs detected by each lab for the indicated protocol. The probability of each miRNA being detected was estimated for every library randomly downsampled to 10^4^, 10^4.5^, 10^5^, or 10^5.5^ total read counts. Simulations and probability estimates were performed using the *drarefy* function, available in the R package, Vegan. A miRNA was only counted as detected if the probability of detection in all technical replicates of a given lab/method was at least 90%. **Note:** Plasma library data was contributed by Labs 1-6. All replicates from NEBNext Lab3 and 4N_Xu Lab5, as well as one replicate from TruSeq Lab1, were excluded from these analyses because of inadequate read counts.

Next we evaluated the reproducibility of plasma sequencing between labs by examining data generated by multiple groups using the TruSeq protocol. Results obtained with the same protocol across labs clustered together in the heatmap representation of all the data (Figure 5a). The median variability across lab measured using %CV was < 33.17% (top/bottom 2%: 11.54, 101.84; n = 8) and measured using QCD was 0.19 (top/bottom 2%: 0.04, 0.60; n = 8) (Figure 5c). The reproducibility of small RNA-seq using RNA isolated from biological samples was therefore comparable to that observed using the synthetic reference RNA samples above.

In order to study potential differences between protocols with respect to the diversity of microRNA sequences recovered from the standard reference plasma RNA, we performed an analysis of the number of miRNAs detected by each protocol, in which we aggregated in-house 4N-type protocols into one group for the sake of comparison. We carried out this analysis using downsampling of datasets so that the same number of total mature miRNA-mapping reads could be compared across protocols, at varying sequencing depths. We found protocols using randomized end-sequence adapters recovered a significantly larger number of microRNAs than those using defined adapter sequences (Figure 5d). In addition, an indirect assessment of miRNA diversity (i.e., % of total reads accounted for by the 10 most abundant miRNAs) was consistent with the conclusion that 4N protocols generate a more diverse profile of microRNAs (***Supplementary Figure 6***).

### Testing a computational strategy for making data comparable across protocols by calculating protocol-specific bias correction factors

The protocol-specific biases highlighted by our analysis of the equimolar synthetic RNA pool sequencing data represent a significant potential barrier to cross-platform data comparison and integration, especially for applications going beyond relative quantification between samples. However, we observed that the sequence-specific bias is reproducible within a given protocol and across laboratories. Given this, we investigated whether datasets from equimolar synthetic RNA pool sequencing using different protocols could be utilized to calculate between-protocol correction factors for each of the human miRNAs in the synthetic RNA pool. We hypothesized that such correction factors could then be applied to datasets generated using differing library preparation protocols, to create transformed data which could be directly compared for the miRNA species present in the synthetic RNA set.

To generate protocol-specific bias factors across several protocols, it is useful to define one of the protocols as a reference, providing a common comparator to which datasets from other protocols are each corrected to. Here we chose to use our in-house 4N libraries as the reference since these exhibited the least bias and are therefore expected to be closest to ground truth. In order to test this correction factor approach, we chose the TruSeq as the library prep method for which data is to be transformed since as it was the common protocol of the study and we had more sequencing results available. To generate correction factors, we then performed differential expression analysis (i.e., 4N in-house libraries data as a reference *vs* data from TruSeq libraries) using the synthetic equimolar pool sequencing datasets, which contain a large number of mature microRNA sequences. Correction factors were calculated for each microRNA based on the difference between its average normalized abundance in datasets from 4N libraries vs. TruSeq libraries. Key assumptions used in these calculations were: that the median mapped read level calculated for a given protocol should match the median for the 4N results given the same RNA input; that the comparisons of the results from a given protocol and the 4N protocol are performed on data processed in the same way that biological samples are processed (i.e., using the exceRpt pipeline and its mapped read outputs). Full details of the calculation and application of correction factors are provided in the Inter-protocol Bias Correction Factors sections of the Computational Methods. Unsupervised hierarchical clustering revealed that the transformed TruSeq samples were more similar to the in-house 4N reference samples than they were to the original untransformed TruSeq values, thereby demonstrating the efficacy of the correction factors on the synthetic RNA training set (***Supplementary Figure 7a***). In addition, the number of reads of the corrected TruSeq were significantly closer the 4N protocol dataset than their uncorrected counterparts (Mann-Whitney test; alternative hypothesis: true location shift is greater than 0: p = 3.33 × 10^−140^; difference in location: 1.609; 95% CI:1.480 inf) (***Supplementary Figure 7b***). Since perfect correction would result in identical counts, the probability distribution of the absolute differences in log CPMs between a perfectly-corrected sample and the reference would then simply be fully concentrated at 0. The value of the median of these difference distributions is thus a quantitative indicator of the effectiveness of the correction (***Supplementary Figure 7b***).

Finally, to test whether these correction factors derived using the synthetic equimolar pool data could be used to successfully be applied to small RNA-seq data from biological samples, we applied the correction factors to the TruSeq data generated from standard reference plasma RNA. After applying correction factors, we observed that TruSeq data clustered with 4N data and that the expression levels of the corrected TruSeq libraries were significantly closer to the 4N values than their uncorrected counterparts (alternative hypothesis: true location shift is greater than 0: p-value = 7.00 × 10^−09^; difference in location 0.522; 95% CI: 0.369 inf) (Figure 6a and 6b). These findings confirmed the favorable effect of applying this correction strategy to inter-convert data generated by different small RNA-seq library preparation methods from biological samples. A full list of correction factors is supplied in Supplementary Table 6.

**Figure 6.**
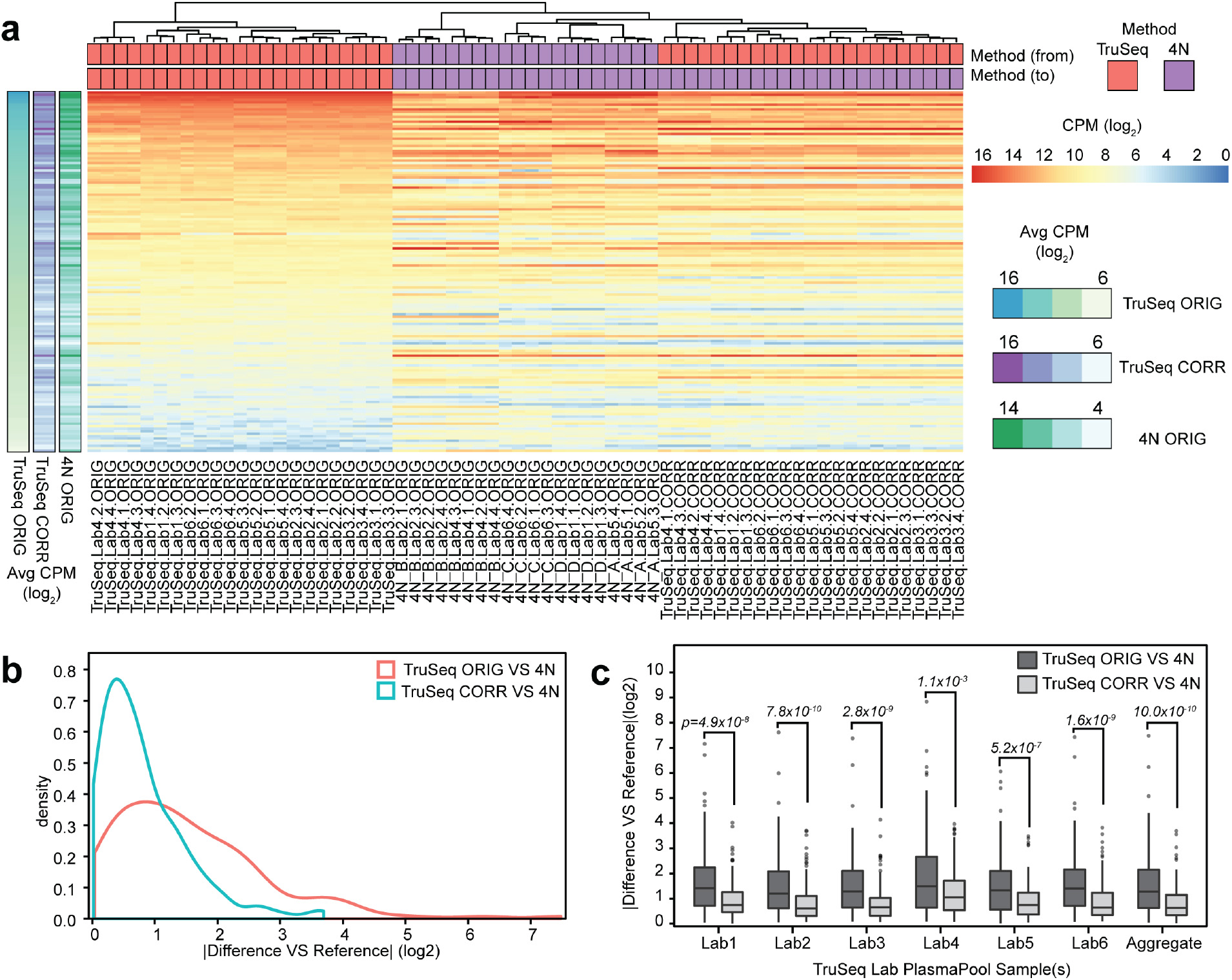
Results of application of computational correction factors to plasma small RNA-seq datasets. (**a**) The heatmap shows mature microRNA CPM (log_2_) values from the reference plasma RNA sequencing datasets corresponding to in-house 4N protocol libraries (4N_A, _B, _C and _D), and those corresponding to TruSeq libraries with and without application of inter-protocol correction factors. Correction factors were calculated using the synthetic equimolar RNA pool data as a training set, and represent the log_2_ fold difference in CPM for each sequence in TruSeq versus 4N libraries. Hierarchical clustering for columns (samples) represents complete linkage clustering on Euclidean distances. Rows (miRNAs) are sorted by mean miRNA abundance in descending order. On the far left, a summary heatmap is shown, depicting the row-wise mean log_2_ expression for 4N plasma libraries, and TruSeq plasma libraries with and without correction. Sample IDs and annotations corresponding to plasma data before or after correction factors were applied are marked as “ORIG” and “CORR”, respectively. Color coding of columns indicates the library preparation method, the four main method groups (TruSeq, NEBNext, CleanTag and 4N). (**b**) Probability density plots show the distribution of average absolute log_2_ fold differences in miRNA CPM between TruSeq and in-house 4N (Reference) plasma libraries before correction (pink line) and after correction (turquoise line) factors were applied. (**c**) Boxplots represent the average absolute log_2_ fold difference in CPM of RNA sequences between uncorrected (dark grey) and corrected (light grey) TruSeq plasma data, relative to the 4N reference, as observed in data from each lab. For each lab, the average log_2_ CPM was calculated for every sequence before and after correction, and the absolute difference was calculated from the average log_2_ CPM measured across all in-house 4N libraries. Wilcoxon rank-sum tests were used to test whether the distribution of the untransformed values versus the 4N reference were significantly higher than that of the transformed values versus reference. P-values for each test are indicated above each plot. The “Aggregate” plot is a boxplot representation and test corresponding to the aggregate data shown in 6b.

### Discussion

Striking differences among results from small RNA-seq experiments have been reported by many researchers. Significant sequence-specific bias has been reported and largely attributed to library preparation protocols^4, 40–42^. This is only one potential source of bias, however, albeit one that is amenable to systematic study. Other sources include more subtle, and difficult to characterize differences that originate in variations in individual laboratory environments and practices. Small RNA-seq differences have not been characterized systematically across multiple labs using different protocols and with common synthetic reference reagents. In a systematic multi-site, multi-protocol study, we designed and profiled three well-characterized synthetic reference RNA sample pools with known amounts of RNAs and a biological plasma RNA pool analyzed in common to evaluate the accuracy, reproducibility, and bias of small RNA-seq methods. The results presented here thus provide a thorough evaluation of the performance of small RNA-seq methodologies, focusing on the potential of using RNA-seq as a quantitative platform. Given the rapid expansion of the use of small RNA-seq in the last few years along with the diversity of protocols, our study comprises an effort towards understanding protocol-specific effects and provides an essential step towards systematic standardization.

Our results have confirmed in a quantitative fashion that small RNA-seq is highly affected by sequence-related bias^4, 40–42, 45^ which is protocol-dependent. The observed biases are as large as 10^4^-fold with some of the commonly used commercial library preparation protocols. This sequence-dependent bias is stronger than the biases previously reported for long RNA-seq^31^, thus highlighting important differences between the technologies and the challenges involved in small RNA and long RNA sequencing. Biases can be particularly vexing when working with low RNA input samples such as biofluids for exRNA analysis, where some low abundance small RNAs may not be reliably detected due to sequence dependent bias.

It is important to note that the in-house protocols used here, which employ adapters containing degenerate bases, reduced the bias on the order of 100-fold. Overall, 4N protocols provided the strongest reduction in the biases. There were, however, differences between the results of 4N protocols using the same adapters. This indicates that there are additional factors in the protocols that affect bias, for example the concentration of PEG in the ligation reactions, time of ligation, etc. Even in the best case with 4N protocols, however, there is still considerable sequence-related bias. This precludes the use of read counts alone for accurate estimation of absolute quantities of diverse small RNAs in a given sample. However, despite the observed biases, we demonstrate clearly here that small RNA-seq is remarkably accurate for relative quantification of small RNA levels, as long as the two samples being compared were subjected to small RNA-seq using the same protocols. In this sense, all of the evaluated protocols were able to distinguish samples with as little as 1.5-fold difference in relative abundance of many of the small RNA species examined. This result thus confirms the suitable role of small RNA-seq as a platform for biomarker discovery and other uses that require only relative measurements.

Reproducibility across laboratories is a crucial requirement for any such experimental method for research and clinical applications^46, 47^. As this can only be tested in extensive comparisons of different sites using standardized protocols and common RNA reference samples, the published small RNA-seq studies to date lack a comprehensive evaluation of reproducibility. To address this issue, we directly evaluated the reproducibility of small RNA-seq both within and between labs in this study. We found that for common protocols small-RNA seq results are remarkably reproducible between labs (%CV ≤ 20% for most of the sequences). This figure is comparable to the %CV reported for “gold standard” techniques such as RT-qPCR^48, 49^. Moreover, when comparing relative quantification results obtained by small RNA-seq across institutions the results were strongly correlated even when the centers were using different protocols for relative quantification. Previous work in which RNA isolation was done separately in each lab^30^ did not report the high degree of reproducibility we observed, and some of the variation in that study is likely to be attributable to RNA isolation-purification.

The use of synthetic RNAs allowed us uniquely to evaluate bias and accuracy because the “ground truth” was known. Since biological material, with a wide range of RNA and other differences could potentially behave differently from the synthetic RNA reference material, we also characterized reproducibility and diversity of miRNAs captured in a common biological sample. The use of standard reference biologically-derived RNA isolated from a pool of human plasma in one center allowed us to gauge the range of detectability of RNA sequences by each protocol. We found that the in-house 4N protocols detected a greater diversity of sequences than those protocols using defined-sequence adapters. It is remarkable that for a given protocol the profile obtained for the biological sample was very reproducible between labs.

The extensive, multi-protocol data generated using a common reference synthetic RNA pool enabled us to compute correction factors for a large number of microRNAs, to convert between datasets generated using different library preparation protocols (e.g., TruSeq to 4N). An inherent limitation of this procedure is that no correction was possible for a small subset of microRNAs that were not detected using both library preparation methods. The correction factor process described here can nonetheless provide an effective method for comparing data obtained using different protocols for many microRNAs. In particular, the equimolar synthetic RNA reference pool described here provides a ready reagent that could be used by investigators to generate protocol-specific correction factors for any protocols, including those that may be developed in the future. Note however, that any change in the adaptor sequence or any change in the reaction conditions would require a new set of correction factors be generated. This method could, of course, be extended to any type of RNA species using synthetic forms of the RNA of interest, and potentially using standardized biological samples as well.

The use of a synthetic equimolar RNA pool allows us to arrange a known, uniform concentration for the synthetic RNAs that is convenient for generating accurate correction factors. In principle, however, correction factors could be obtained using biological RNA samples as long as exactly the same RNA sample was used for cross-protocol comparisons, though some RNAs present at very low abundance in biological material may be subject to greater variation. In addition, the synthetic RNA reference data generated here can enable development of more advanced computational approaches for cross-platform data normalization, beyond the first-generation of inter-protocol correction factors described here. This could include bias corrections within individual library datasets to better estimate the relative abundance of different RNAs within the same sample, which would advance small RNAseq closer to providing information on absolute quantities of small RNAs in a sample.

In summary, the results of this study demonstrate both the limitations and the significant potential of small RNA-seq as a quantitative platform for multi-institutional studies of small RNA biology and biomarker discovery. As new modifications of the technology are developed, we suggest that the kind of cross-platform, cross-laboratory study described here will provide a model and reference data for characterization of additional methods. While there is certainly room for improvement of the technology for small RNA measurement by RNA-seq, our study confirms the reproducibility of small RNA-seq when specific, standardized protocols are carefully followed. It also demonstrates the value of the approach for relative change studies, and the possibility of increasing the cross-platform comparability of RNA level measurements using carefully obtained correction factors.

## Acknowledgements

We acknowledge funding support from the NIH Extracellular RNA Communication Common Fund grants: U01 grants HL126499 to M.T., HL126496 to D.G and K.W., HL126493 to D.J.E. and P.G.W., HL126494 to L.C.L., HL126495 to J.E.F., HL126497 to I. G., and UH3 grant TR000891 to K.V.-K-J. M.D.G. acknowledges initial support from a Rio Hortega Fellowship and later from a Martin Escudero Fellowship. E.N.M.N-TH. and T.A.P.D. received funding from the European Research Council under the European Union’s Seventh Framework Programme (FP/2007-2013) / ERC Grant Agreement number 337581. Y.W. received funding from the Dana-Farber Strategic Plan Initiative. K.W. received funding from DOD (W911NF-10-2-0111) and DTRA (HDTRA1-13-C-0055). Research reported in this publication was also supported by the National Cancer Institute of the National Institutes of Health under Award Number P30CA046592 by the use of the following Cancer Center Shared Resource at the University of Michigan: DNA Sequencing. The content is solely the responsibility of the authors and does not necessarily represent the official views of the National Institutes of Health. We are grateful to Joel S. Rozowsky, Robert Kitchen, Sai Lakshmi Subramanian, William Thistlethwaite, Mark B. Gerstein, Aleksandar Milosavljevic and Nikita Sakhanenko for facilitating access to the exceRpt pipeline and helpful conversations and suggestions.

## Author Contributions

M.D.G. designed, led and coordinated the overall study. This comprised contributions throughout the entire process including designing, preparing and distributing synthetic RNA pools, creating detailed instructions for the participating labs, performing experiments, coordinating experimental work and communication across labs, and organizing and managing all aspects of the project. In addition, she interpreted study results and was the primary writer of the manuscript.

R.S. led the computational analyses and data management for this study, including processing data, designing and performing data analyses, identifying and applying methods to visualize data and results, and coordinating data and metadata incoming from collaborating laboratories. He also designed the composition of the ratiometric pools, interpreted results and contributed to the manuscript by preparing the figures, drafting figure legends and writing the computational methods.

A.E. contributed to experimental design and preparation of synthetic RNA pools. He also performed experimental work, interpreted results and provided comments on the manuscript. He also developed and contributed the core “in-house” 4N protocol, variations of which were then utilized by multiple laboratories.

M.T., D.G. and D.J.E. helped design the study and interpreted results, along with contributions from the rest of the study team including in particular L.L.C., P.G.W., K.V.K-J., I.G., Y.W., K.W., J.E.F., and E.N.M.N.H. H.J. contributed to design of statistical analyses and data interpretation. P.M.G., A.J.B., S.S., P.L.D.H, K.T., A.C., L.S., J.K., R.R., D.B., and T.A.P.D. carried out experiments. M.T. and D.G. supervised the overall study and did primary editing of the manuscript with significant input from D.J.E. However all authors contributed to reviewing, editing and/or providing comments on the manuscript.

## Online methods

### Experimental methods

#### Reference samples

A synthetic equimolar pool containing 1,152 synthetic RNA oligos was prepared in an RNase-free environment and working on ice to minimize degradation. The pool was prepared by combining (i) the miRXplore Universal Reference from Miltenyi Biotec Inc (Auburn, CA), which comprises 962 RNA oligonucleotides with sequences matching human and other microRNAs, and (ii) a set of 190 additional, custom-synthesized RNA oligonucleotides, to generate the pool in which each of the 1,152 RNA oligonucleotides is present at equimolar concentration. This latter set comprises microRNAs and non-microRNA sequences of varied sequence and length from 15 to 90 nt, which were synthesized, HPLC-purified and quantified spectrophotometrically by IDT, Inc. (Coralville, IA). The resulting equimolar pool was aliquoted in prelabeled DNA-, DNase-, RNase-, and pyrogen-free screw cap tubes with low adhesion surface and stored immediately at −80C. Aliquots were distributed to the participant laboratories in overnight shipments with an abundant supply of dry ice. The complete list of RNA sequences comprising the equimolar pool is provided in **Supplementary Table 1**.

Two ratiometric pools, SynthA and SynthB, containing 334 synthetic RNA oligonucleotides also synthesized by IDT (in some cases comprising microRNA and non-microRNA sequences from the set of 190 described above), of which subsets of sequences were present in 15 different varying ratios between the two pools were designed in the coordinating lab (see computational methods). These were also prepared, aliquoted and distributed to the participant centers following the same previously mentioned precautions to avoid RNA degradation. The complete list of sequences comprised in the ratiometric pools as well as their different ratios in SynthA and SynthB are provided in **Supplementary Table 2**.

Plasma samples from eleven healthy male donors with age ranging from 21-45 years were obtained and pooled in one of the participating labs (***Supplementary protocol 1***). RNA from this pool of plasma was isolated (***Supplementary protocol 2***), mixed and distributed to the rest of participant centers.

#### Library preparation and small RNA-seq

A written guideline for library preparation and sequencing was distributed to all the participant centers. The input RNA for library preparation was 10 femtomoles for the synthetic pools and 2.1 ul for the plasma RNA pool. Each group prepared four replicate libraries from each sample using the following small RNA library preparation protocols (the labs marked with an asterisk did not contribute plasma libraries): Lab1 (TruSeq and in-house 4N_D), Lab2 (TruSeq and in-house 4N_B), Lab3 (TruSeq and NEBNext), Lab4 (TruSeq, in-house 4N_B and NEBNext), Lab5 (TruSeq, CleanTag and in-house 4N_A and 4N_Xu), Lab6 (TruSeq and in-house 4N_C), Lab 7* (NEXTflex), Lab 8* (TruSeq and NEXTflex) and Lab 9* (TruSeq and NEBNext).

The protocols for TruSeq, CleanTag, NEBNext and NEXTFlex for Illumina were performed according to the manufacturer’s instructions (note that for NEBNext, adapters were diluted, 1:2 in Lab3 and Lab9 and 1:6 in Lab4. For CleanTag 1:20 dilution of the adapters was performed). NEXTflex for Ion Torrent sequencing was performed as described in ***Supplementary protocol 3***. In-house 4N protocols A, B, C and D were performed as described in ***Supplementary protocol 4-7***. 4N_Xu protocol was performed as previously described48 50. Size selection was performed using Pippin Prep (in Lab1, Lab2, Lab4 and Lab6 for all the protocols), TBE gels (in Lab3, Lab5 and Lab9 for all the protocols and Lab 8 for TruSeq) or beads (in Lab7 and Lab8 for NEXTflex).

Single-end libraries were sequenced using the Illumina HiSeq 2500 (Lab5, Lab8 and Lab9), Illumina HiSeq 4000 (Lab1 and Lab4), Illumina NextSeq 500 (Lab2, Lab3 and Lab6) or Ion Torrent (Lab7) platforms. All labs using Illumina platform sequencers generated 50 bp single-end reads, except for Lab9, which generated 75nt single-end reads. Each laboratory was free to choose the number of samples to pool per lane but a target of at least 8 million reads per library was requested. FASTQ files were uploaded to the Genboree workbench for central data analysis.

### Computational methods

#### Designing Composition of Ratiometric Pools

290 artificial sequences were assigned at random to 8 ratiometric groups (1, 1.5, 2, 3, 4, 5, 8 and 10x) and to either ratiometric SynthA or SynthB. The ratio indicates how much more concentrated the sequence would be in the assigned sample than the other. In other words, a sequence in the 10x pool assigned to SynthA would be pooled at the base concentration in SynthB and at 10 times the base concentration in SynthA. To make groups of approximately equal size, we took 8 sequences at a time and assigned them to the 8 ratiometric groups randomly, without replacement. To ensure the total amount of oligonucleotide was approximately equal in sample A and sample B, an even number of sequences was assigned to each ratiometric group and were distributed equally between sample A and B, using a similar method of equally-distributed random assignment. The random assignment was performed in Excel and the results are shown in **Supplemental Table 2**.

#### NGS barcode splitting, FASTQ generation and data coordination

High-throughput sequencing, barcode splitting and FASTQ file generation was performed by each of the participating groups independently. FASTQ files were uploaded to the Genboree Workbench for centralized read preprocessing, mapping, read categorization and counting using the ExceRpt small RNA analysis pipeline. (http://genboree.org/java-bin/workbench.jsp).

#### Preprocessing, mapping and read counting

FASTQ files for the Equimolar, Ratiometric and Plasma pools were initially processed through the exceRpt small RNA-seq Pipeline (Version 4.3.5), using the batch submission tool. For details on the exceRpt pipeline and the associated processing steps, see the Genboree Workbench documentation (http://genboree.org/theCommons/projects/exrna-tools-may2014/wiki/Small%20RNA-seq%20Pipeline). A brief description of parameters that were changed from the default, or that differed between libraries, is included below.

The exceRpt pipeline was used at the default settings as much as possible. The default setting for adapter trimming is to “auto-detect” and trim the adapter found in each file, and all samples were initially submitted using this functionality. Auto-detection was also used for the 4N libraries, which contain 4nt random sequences at the 5’ and 3’ ends of the insert sequence; however, the length and position of the random bases was manually entered. The sequence and identity of the adapter identified by the exceRpt pipeline was confirmed in the output files. Any library with a missing or incorrect adapter identified was re-submitted to the exceRpt pipeline, and the correct adapter sequence manually entered.

By default, sequences less than 18 nt after adapter trimming are removed and not used for downstream analysis. This default minimum read length was used for Plasma Pool libraries. The minimum read length for the synthetic pools was changed to 15, which corresponds to the length of the shortest sequences in the Equimolar and Ratiometric Pools.

Equimolar, Ratiometric SynthA and Ratiometric SynthB libraries were mapped to a “Spike-In” sequence library uploaded to the Genboree Workbench. This file is in FASTA-format and contains a non-redundant set of sequences from the Ratiometric and Equimolar Pools (***Supplementary Table 7***). When a Spike-In file is provided to the exceRpt pipeline, a bowtie2 index is created. Adapter-trimmed and filtered reads are mapped to the spike-in index with bowtie2. The number of reads aligning to each sequence is included in the output, and was used for all downstream analysis for the synthetic pools.

Plasma pool libraries were mapped to hg19 using the STAR alignment algorithm (the default option for exceRpt). Multi-mapping-adjusted read counts from reads mapping to mature miRNAs is provided in the exceRpt pipeline output, and was used for downstream analysis and comparison of the plasma pool libraries.

##### Sample Filtering

Unless specifically noted in the text, libraries were excluded from the analysis whenever a minimum mapped read count threshold was violated. For the synthetic pools, an average of one million reads mapping to the “spike-in” sequences were required across all replicate libraries. For the plasma pool samples, replicate libraries with fewer than 100,000 miRNA-mapping reads were removed. The entire sample was removed if more than one of the replicate libraries failed to pass the minimum count threshold.

##### Equimolar Pools Analysis

Read counts for the equimolar (and likewise for the ratiometric pools) were obtained from “calibratormapped.counts” files included in the exceRpt pipeline output for each sample file. Sample-specific information, including the contributing lab, library preparation method and replicate number were associated with the corresponding calibrator count file, and were loaded into R for analysis. A full list of equimolar and ratiometric sequences with additional sequence information was used as a reference to merge counts from all input files and add in zero counts, where needed. Unless specifically mentioned in the text, analysis of ratiometric and equimolar libraries was limited to sequences with a 5’-Phosphate modification, 16-25nt in length. Scaling read counts to counts-per-million was calculated using the total counts from the filtered sequences. For any measurements or comparisons of CPM across labs, CPM values were normalized between labs using the “relative log expression” normalization implemented in the R package, edgeR (calcNormFactors; method=”RLE”).

##### Determining Over-represented and Under-represented Sequences

Equimolar sequences in each replicate library were ranked by abundance, assigning the minimum rank value in case of ties. The 10 top and bottom-ranked sequences were determined by assessing ranks from counts arranged in descending and ascending order, respectively. TruSeq, NEBNext, CleanTag and 4N libraries were each queried for sequences consistently found in the top or bottom 10, as defined by at least 75% agreement among the libraries of at least one method.

##### Ratiometric Pools Analysis

Ratiometric pool counts were initially processed along with the equimolar pool counts, as described above, keeping read counts for 16-25 nt sequences present in the ratiometric pool input library. The ratio of SynthA: SynthB was calculated as the ratio of the mean CPM across technical replicates in SynthA / SynthB. If the mean CPM in SynthA and SynthB were both 0, the ratio was set to 0.

###### Ratiometric Pools: Differential Expression

Independent differential expression workflows were run for each lab and library prep method, following a standard two-group comparison between “A” and “B” ratiometric pools. Normalization, dispersion estimation and differential expression testing was performed using three different R packages: *EdgeR*, *DESeq2* and *limma/voom*. For *EdgeR*, dispersion estimates were calculated using the Relative Log Expresion (RLE) method, and significance was calculated based on a likelihood ratio test on the null hypothesis that ratiometric sample B − A = 0. Default settings were used for *DESeq2*, and significance was calculated based on a Wald Test. Significance for the *limma/voom* workflow was based on and empirical Bayes moderated t-test.

#### Plasma Pools Analysis

Comparison of plasma pool libraries was limited to mature miRNAs. Read counts for mature miRNAs were taken from “readCounts_miRNAmature_sense.txt” files provided in the exceRpt pipeline output. The read count files and associated metadata for all samples were loaded and merged in R for further analysis. Multi-mapping-adjusted read counts are calculated as part of the exceRpt pipeline and were used for all comparisons. The total number of unique reads mapping to miRNAs was taken from “.stats” files provided in the exceRpt pipeline output.

#### Downsampling Read Counts

The R package, Vegan, was used to simulate random downsampling of Equimolar and Plasma pool count matrices. The Vegan function, *drarefy*, was used to estimate the probability of detection for each sequence based on random simulations of downsampling to specified levels. For the plasma pools, downsampling was performed to four different levels (10^4^, 10^4.5^, 10^5^ and 10^5.5^). Equimolar pools were downsampled to six different levels (10^6.5^ down to 10^4^ at half-log intervals). Libraries with read counts below the specified threshold are removed. The probability of each miRNA being detected in all technical replicates of a given sample was calculated as the product of the probability estimates for the individual replicates.

#### Inter-protocol Bias Correction Factors: Estimation

Equimolar pool samples were processed through the exceRpt pipeline using the same input parameters as the plasma pool libraries, in order to obtain multi-mapping, scaled read counts for mature miRNAs that were directly comparable to the plasma pool counts. Differential expression analyses were performed using the mature miRNA read counts for the equimolar pool samples, and scaling factors were taken from the resulting log_2_ fold-change estimates. For details on *limma* and *voom* functionality and the parameters used, see the documentation for the *limma* package.

To summarize: scaling factors were calculated for each pair of library prep. methods using the following workflow:

1. Filter out miRNAs with 0 counts in any library: For the subset of samples being tested, scaling factors are only calculated for miRNAs having at least one count in every sample of the two methods being tested.
2. Prepare miRNA count matrices for linear modeling: Use the R package, *voom*, to calculate precision weight estimates and normalize data to allow count data to be analyzed appropriately using the limma package. Normalization is also performed between samples such that the median miRNA expression value is the same in all samples.
3. Fit miRNA-wise linear models to account for batch (lab) effect: The lmFit function from the *limma* package is used to fit linear models for each miRNA, estimating coefficients for each lab + library prep method. The coefficients represent the differences in expression for each miRNA between each lab + library prep method.
4. Estimate the log fold change between the two methods for each miRNA: Use the fitted model to calculate for each miRNA the average expression estimated using the method A coefficients - the average expression estimate for the method B coefficients.

The R packages *limma/voom* were used for read count normalization and differential expression estimates, using standard workflows suggested for RNAseq data to account for batch (lab) effects, and then testing for the main effect of the library prep methods. For each pairwise comparison of library preparation methods, equimolar pool counts matrices were extracted and only miRNAs with >= 1 read count in all samples of both methods were kept for analysis. After filtering, *voom* was used to normalize the count data and calculate precision weight estimates that allow count data to be appropriately tested with the linear modeling schema used in the *limma* package. *Voom* was run with the default parameters, except that read counts were additionally normalized between arrays using the “scale” method, which adjusts read counts such that the median miRNA expression value is the same in all labs. The *voom*-transformed data was supplied to the *limma* lmFit function, along with a design matrix indicating the coefficients to be estimated. Initially, coefficients were estimated for each lab + library prep method in order to model batch/lab-specific effects. The main effect of the library prep method was then calculated as the average effect of method 1 - method 2. A contrasts matrix was generated and supplied, with the fitted model, to the contrasts.fit function, followed by an empirical Bayes function to estimate the resulting statistics for each miRNA. Log_2_ fold-change estimates, along with 95% CI were obtained from these estimates.

#### Inter-protocol Bias Correction Factors: Applying corrections

The equimolar pool-derived, inter-protocol bias correction factors were applied to the corresponding plasma pool samples for testing. To apply the correction factors, count matrices for the subset of plasma pool libraries being compared were selected and were then pre-filtered and normalized in the same way as the equimolar pools in generating the correction factors, described above. MiRNAs were filtered to include only those with a) a correction factor estimated from the equimolar pool and b) at least 5 counts in every library in the subset of methods being compared. Correction factors were applied to the appropriate samples. For example, if correction factors were calculated as the log_2_ fold-change between TruSeq and 4N samples (TruSeq- 4N), then the correction factors would be applied to the log_2_-transformed TruSeq plasma pool samples by subtracting the correction factor. For the heatmaps and density plots, corrected values were added to the original count matrix of untransformed values, and unless specifically noted in the text, normalized using quantile normalization.

## Supplementary figures

**Supplementary Figure 1. Examples of protocol-dependent bias in small RNA-seq.**

Bar plots show the average CPM (y-axis), measured across TruSeq (pink), CleanTag (green), NEBNext (turquoise) and all 4N (purple) libraries, for individual sequences (x-axis) consistently (**a**) over-represented, or (**b**) under-represented, in at least one library preparation protocol. The over- and under-represented sequences plotted were among the 10 highest or 10 lowest-expressed, respectively, in >=75% of TruSeq, CleanTag, NEBNext or 4N libraries. Relevant to both panels in this Figure, over- and under-represented sequences were determined for each protocol (i.e. TruSeq, NEBNext, CleanTag and 4N) separately. Sequences with the 10 highest and 10 lowest CPM values were identified in each of the libraries. Over- and under- represented sequences for a particular library preparation protocol were selected on the basis of being among the top or bottom 10, respectively, in at least 75% of libraries for a given protocol.

**Supplementary Figure 2. Number of sequences with a low probability of detection in the equimolar pool at different sequencing depths.** Bar plots show the number of sequences in the equimolar pool with a probability of detection <90%, as estimated from random simulations of downsampling all libraries to equal levels. Total miRNA-mapped reads were downsampled to six different levels–from 10^6.5^ down to 10^4^ at half-log intervals–as indicated in the grey boxes above each plot. Random simulations and probability estimates were performed on replicate libraries using the *drarefy* function included in the R package, Vegan. The probability of each miRNA being detected in all technical replicates of a given sample was calculated as the product of the probability estimates for the individual replicates. Bars are grouped and color-coded by library preparation method (Truseq – pink; CleanTag – green; NEBNext – turquoise; 4N – purple). The number of equimolar pool sequences with less than 90% probability of detection by all technical replicates are shown above each bar. “NA” labels indicate samples where libraries had total miRNA-mapped read counts than the downsampling level indicated, and therefore were not able to be evaluated at that level.

**Supplementary Figure 3. Spearman correlation for the Equimolar pool.** Heatmap showing the squared Spearman rank correlation coefficients calculated based on mean CPMs for sequences in the equimolar pool. CPMs for each sequence were first averaged across technical replicates, before running pairwise comparisons. Rows and columns are color coded to indicate different labs and library preparation protocols.

**Supplementary Figure 4. Intralab reproducibility of CPM measurements in datasets from sequencing the SynthA and SynthB pools.** Violin plots showing the % Coefficient of Variation (%CV) and quartile coefficient of dispersion (QCD) calculated across technical replicates of the SynthA and SynthB libraries. Plots are grouped and color-coded by library preparation method (Truseq – pink; CleanTag – green; NEBNext = turquoise; 4N = purple), with SynthA and SynthB libraries shown as lighter or darker shades, respectively. Horizontal lines within each violin indicate the 1^st^, 2^nd^ and 3^rd^ quartiles.

**Supplementary Figure 5. Small RNA-seq accuracy in detecting differentially abundant miRNAs between ratiometric synthetic RNA pool samples SynthA and SynthB.** Barplots show the fraction of miRNAs in each ratiometric subpool that were found with significantly different CPM values (adjusted p-values < 0.01) measured between the SynthA and Synth B samples. Tests for significant differences in abundance were performed using three different R packages commonly used for RNA-seq differential expression analysis: DESeq2 (pink), EdgeR (green), and limma/voom (blue) (see Methods for details). P-values are calculated based on the default tests for each of the three algorithms: likelihood ratio test (EdgeR), Wald Test (DESeq2) or empirical Bayes moderated t-test (limma) (H0 = SynthB − SynthA = 0, FDR <= 0.01, n = 4 per sample, except from TruSeq Lab8 SynthB = 3). Each plot represents independent tests of differential expression for each lab + library prep method.

**Supplementary Figure 6. Plasma microRNAs with the highest number of reads.** The bar plot shows the fraction of miRNA-mapping reads corresponding to the 10 miRNAs with the highest read counts sequenced in each sample. The y-axis shows the sum of the multi-mapping-scaled read counts for the top 10 mature miRNAs / sum of the multi-mapping-scaled read counts for all mature miRNAs. The x-axis separates distinct samples, with each bar grouped and different shades of the same color representing different technical replicates. Samples are color-coded by library preparation protocol (TruSeq – pink; CleanTag – green; NEBNext – turquoise, 4N-purple).

**Supplementary Figure 7. Calculation of correction factors in the Equimolar pool. (a**) The heatmap shows expression (quantile-normalized log_2_ CPM) of equimolar pool sequences measured from in-house 4N protocol samples (4N_A, _B, _C and _D — all combined and referred to here as 4N and coded as purple columns), and from TruSeq samples (coded as pink columns) before and after inter-protocol bias correction factors were applied. Correction factors were calculated from equimolar pools to correct TruSeq to 4N methods. Hierarchical clustering for columns (samples) represents complete linkage clustering on Euclidean distances. Rows (miRNAs) are sorted by mean miRNA abundance in descending order. Row-wise mean log_2_ expression for 4N, and TruSeq with and without correction for each row is provided at the far left. (**b**) Probability density plots show the average absolute log_2_ fold difference measured for each sequence between the in-house 4N (Reference) and the TruSeq equimolar pool samples before correction (pink line) and after correction (turquoise line) factors were applied.

## Supplementary tables

**Supplementary Table 1. Synthetic Equimolar pool.** Complete list of sequences comprising the equimolar synthetic pool.

**Supplementary Table 2. Synthetic ratiometric pools (SynthA and Synth B).** Complete list of sequences comprised in the ratiometric pools as well as their different ratios in SynthA and SynthB.

**Supplementary Table 3. Quality control.** Table showing quality control metrics for all the libraries prepared for the study.

**Supplementary Table 4. Number of undetected sequences in the Equimolar pool.** Table showing the total number of reads and the number of undetected sequences (sequences with zero reads in ≥ replicate) per each lab/protocol.

**Supplementary Table 5. Observed *vs*. Expected ratios.** Table listing the number and % of sequences within 1.5 fold, 2-fold, and 2.5 fold of expected per each lab /protocol.

**Supplementary Table 6. Correction Factors.** Table providing the correction factors calculated from the equimolar pool between all protocols. The “logFC” column indicates the correction factor value, and CI95L/R columns provide a 95% CI for the correction factor estimate.

**Supplementary Table 7. Non-redundant set of sequences from the Ratiometric and Equimolar Pools**

**Supplementary protocols.**

**Protocol 1.** Plasma separation from whole blood.

**Protocol 2.** RNA isolation from plasma pool.

**Protocol 3.** NEBNext small RNA library preparation for Ion Torrent sequencing platform.

**Protocol 4.** In-house 4N protocol A

**Protocol 5.** In-house 4N protocol B

**Protocol 6.** In-house 4N protocol C

**Protocol 7.** In-house 4N protocol D.

